# Threshold responses of aquatic microbial diversity to terrestrial land use across Mediterranean catchments

**DOI:** 10.64898/2026.08.07.743446

**Authors:** Carmen Soler-Zamora, Emilio Cano, Patrizia Elena Vannucchi, Enrique Lara, Bertrand Fournier

## Abstract

Climate driven aridification and intensified human activity are placing increasing pressure on Mediterranean freshwater ecosystems. These impacts propagate from land to water, altering nutrient regimes and reshaping aquatic microbial communities. We analysed Arcellinida diversity across 363 lentic inland saline and freshwater sediment samples spanning broad gradients of land use, water chemistry, soil properties, and climate in southern Spain. Random forest models identified terrestrial land use intensity followed by water chemistry as main predictors of community diversity. Diversity declined sharply in sites with population densities above ∼33 inhabitants/km² and under eutrophic conditions, but peaked in oligotrophic systems with stable, carbon rich soils. These threshold responses demonstrate that aquatic protist assemblages integrate both long term terrestrial pressures and current water conditions. Overall, our findings show that landscape transformation and its cascading effects on water quality dominate community assembly, and that the combination of community level diversity metrics with selected taxon-level indicators capture ecosystem degradation more consistently than relying on a single metric.

## 1. Introduction

Freshwater ecosystems occupy less than one percent of Earth’s surface, yet they sustain a disproportionate share of global biodiversity and ecosystem services. They regulate floods, recharge groundwater, support agriculture, and harbour nearly ten percent of all known animal species (Lynch et al., 2023; Sayer et al., 2025). As freshwater ecosystems are embedded within terrestrial landscapes, their condition and diversity are inseparable from the land that drains into them. Nutrient and sediment fluxes link land, water, atmosphere, and human activities on land—agriculture, urbanization, industrialization—propagate through these connections to alter freshwater chemistry, productivity, and biotic structure (Albert et al., 2021; Carpenter et al., 1998; McFadden et al., 2023). These interactions are more pronounced in a region settled since millennia as the Mediterranean basin, where intensified land use and climate-induced aridification jointly threaten freshwater ecosystems integrity (Estrela-Segrelles et al., 2023; Strayer and Dudgeon, 2010).

Understanding biodiversity alterations in such coupled systems requires integrating both aquatic conditions and the slower terrestrial processes that shape them. Although water chemistry can fluctuate over days to weeks, land use patterns often persist for decades, leaving long-term imprints on ecological communities. Organisms differ in how quickly they respond to these drivers: bacterial assemblages can reorganize within days in response to nutrient pulses (Sun et al., 2017), whereas longer-lived or dispersal-limited taxa such as fish or benthic invertebrates integrate environmental conditions over years (Allan and Flecker, 1993; Townsend et al., 2008). High-resilience communities therefore record the cumulative effects of land use change, soil degradation, and climate forcing, rather than the transient hydrological states captured by short-lived taxa (Flinn and Vellend, 2005; Haddad et al., 2015; Peay et al., 2010).

Protists occupy a pivotal position in aquatic food webs, linking primary production, decomposition, and predation (Bjorbækmo et al., 2020; Guo et al., 2023). Their community structure reflects both bottom-up nutrient supply and top-down control, making them sensitive indicators of ecosystem-level change (Payne, 2013). Compared with prokaryotes, microbial eukaryotes often respond to longer-term or spatially integrated environmental gradients; for example, protist diversity in riparian zones correlates more strongly with local habitat conditions than with short-term seasonal shifts (Fournier et al., 2020).

Among protists, Arcellinida (lobose testate amoebae) are particularly suitable for exploring the coupling between terrestrial and aquatic systems. These organisms are particularly abundant and diversified in lakes and ponds, where they have been studied since early 20^th^ century (Penard, 1902). They are microbial top predators, and depend therefore on healthy food webs to thrive (González-Miguéns et al., 2024). Arcellinida are considered as K-strategists among protists; they have slow growth, occupy narrow ecological niches, and are sensitive to multiple stressors including nutrient enrichment, heavy metal pollution, salinization, and hydrological disturbance (Charqueño Celis et al., 2019; Escobar et al., 2008; Kosakyan and Lara, 2019; Nasser et al., 2020; Patterson et al., 2013; Reinhardt et al., 1998; Roe et al., 2010). Their slow population turnover and capacity for dormancy allow communities to integrate environmental conditions over extended periods, making them natural recorders of long-term landscape influence.

This research investigates how terrestrial land use, in-water chemistry, soil properties and climate jointly shape the diversity and composition of Arcellinida diversity in water bodies (reservoirs, lakes and slow-flowing rivers) across Southern Spain. We test two hypotheses: (1) that Arcellinida diversity primarily reflects slow-changing processes, particularly land use, in line with their K-strategist lifestyle; and (2) that increasing human land use intensity is associated with lower diversity and changes in community composition. Through this lens, we assess how the transformation of Mediterranean landscapes under global change governs the organization of aquatic microbial life.

## 2. Material and methods

### 2.1. Study region and sampling design

We investigated the effects of terrestrial land use on freshwater microbial diversity across 363 samples from 147 sites distributed throughout southern Spain (36–39°N, 1–8°W) (Fig. 1). Sampling sites included lentic inland saline and freshwater bodies from reservoirs, lakes and slow-flowing rivers spanning a gradient of surrounding land use from natural forest to intensive agriculture and urban areas. We collected superficial sediment samples during two campaigns in 2023, in spring (maximum water level) and in summer (minimum water level). This sampling program aimed to capture key seasonal variability and, therefore, provide a realistic picture of Arcellinida communities all year round.

**Fig. 1.**
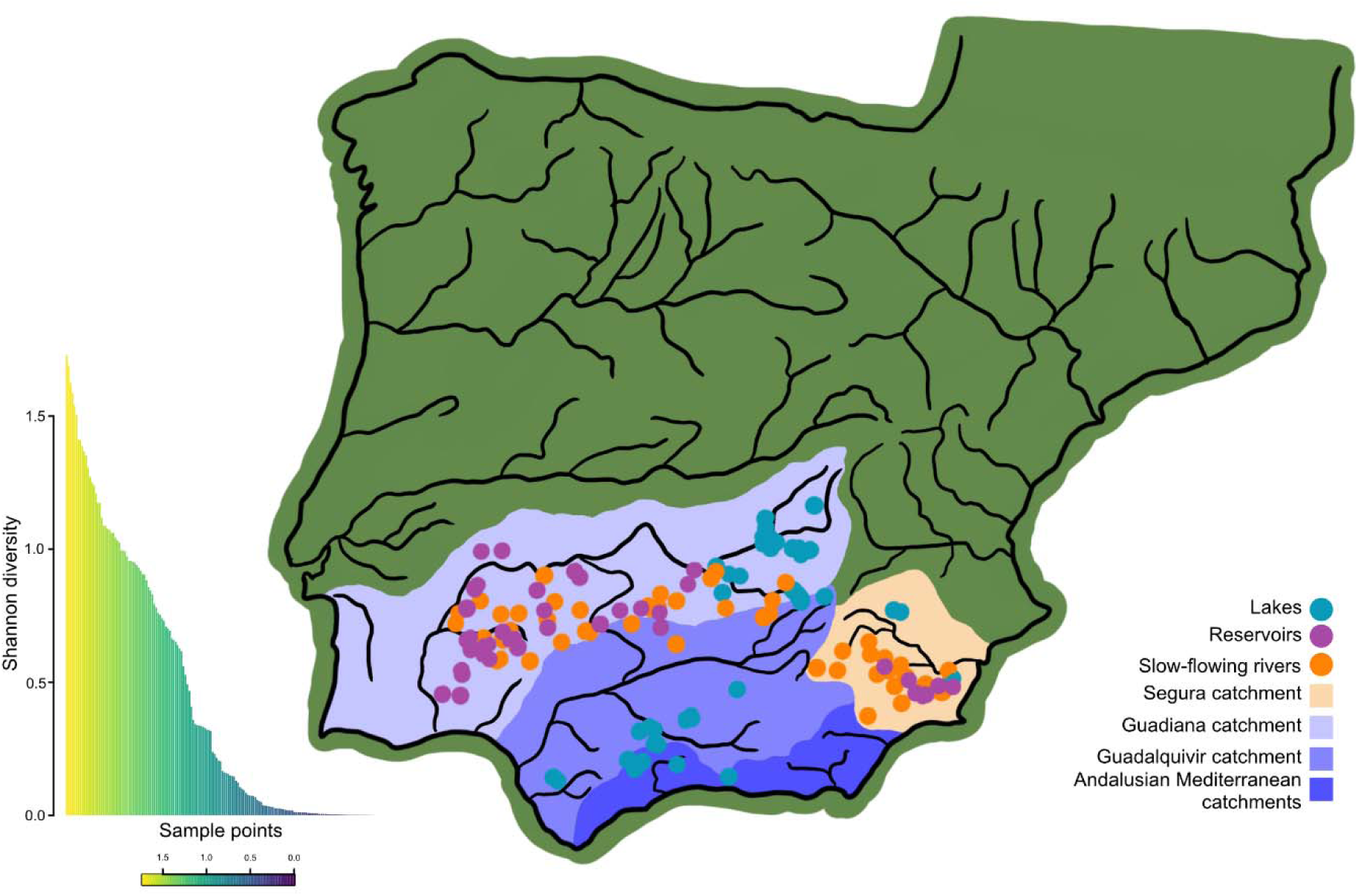
Study area characterization and Arcellinida diversity. (a) Geographic distribution of sampling sites across catchments. Circles indicate waterbody types: lakes (blue), reservoirs (purple), and slow-flowing rivers (orange); colours delineate individual catchments. Map designed entirely by Carmen Soler-Zamora. (b) Shannon diversity indices calculated from OTU data for each sampling site. Bars represent individual samples, ordered from highest (yellow) to lowest (dark blue) diversity.

At each site, we collected two replicates of top-most few millilitres of sediment (∼1 mL) using sterile disposable plastic Pasteur pipettes. Samples were immediately preserved in a 1:1 volume of LifeGuard Soil Preservation Solution (Qiagen) and stored in sterile 2 mL tubes to avoid cross-contamination.

### 2.2 eDNA extraction and amplification

We extracted total eDNA using the DNeasy PowerSoil Pro Kit (Qiagen) following the instructions provided by the manufacturer and preserved frozen (−20°C) until further processing.

We used a two-step nested polymerase chain reaction (PCR) protocol specifically designed for Arcellinida to amplify a fragment of 640 bp from the cytochrome oxidase subunit I (COI). We performed the first PCR with Folmer’s primers (Folmer et al., 1994) and second with a specific primer, as described elsewhere (González-Miguéns et al., 2024, 2023; González-Miguéns et al., 2022) but using Invitrogen™ Platinum™ SuperFi™ DNA Polymerase. For the second PCR, we used a unique combination of primers with tags for each sample (Table S1) amplifying a fragment of 350 bp as described by González-Miguéns et al. (2023).

Finally, we quantified the eDNA using a Qubit 3 fluorometer (Invitrogen), with dsDNA high-sensitivity (HS) assay kits (Thermo Fisher). We normalized the DNA concentration of all samples and generated two pools with the resulting amplicons. The Genomic Unit of the Fundación Parque Científico de Madrid (Spain) sequenced these pools in one Illumina NextSeq P1 600 cycles (2×300) run.

### 2.3. Data curation and “tag-jumping”

The curation of the sequences implied: a) trimming of primers and demultiplexing using cutadapt v2.8 following the pipeline “Cutadapt_pipeline.bash”. b) The resulting reads per sample were analysed with the DADA2 R package (Callahan et al., 2016) following the tutorial in https://benjjneb.github.io/dada2/tutorial_1_8.html, with minor modifications, after (González-Miguéns et al., 2023). This procedure provided Amplicon Sequence Variants (ASV). c) We used VSEARCH v 2.14 (Rognes et al., 2016) for the taxonomic assignment of the ASVs using eKOI, a COI curated database built on publicly available sequences with a special emphasis on protists (González-Miguéns et al., 2025).

Samples were bioinformatically sorted using sample-specific indexes (“tags”), which implies a risk of sample misassignment, as fluorescent signals can overlap in the sequencing by synthesis process (“tag jumping”) (Rodriguez-Martinez et al., 2023; Schnell et al., 2015). To limit this bias as much as possible, we carried out a combination of the protocols developed in earlier works (ElKhouri-Vidarte et al., 2025; González-Miguéns et al., 2023; Useros et al., 2024) to mitigate the tag jumping biases in results interpretation. We only brought minor modifications, as explained below.

The necessary thresholds to remove misassigned sequences were determined using the same approach as in (ElKhouri-Vidarte et al., 2025); the idea behind this approach is that, for eliminating ASVs as probable tag jumping artefacts, we determined thresholds by observing ASVs that appeared in unused tag combinations (=“blank samples”). In a first step, we eliminated all ASV below a minimum read count of seven, thus excluding those with too few reads that might be associated with errors or noise. Then, we calculated the thresholds based on read proportions between “blank” and “real” samples.

In addition to the protocols developed in (ElKhouri-Vidarte et al., 2025; González-Miguéns et al., 2023; Useros et al., 2024), we discarded sequences that did not have the expected length because COI is a protein-coding gene. As 350 bp is the optimum length for ASV, we filtered out sequences ranging between 345 to 365 characters. Finally, we selected only those ASVs that had an identity value for Arcellinida higher than 84% for the downstream analyses; this threshold having been determined empirically (González-Miguéns et al., 2024).

### 2.4. Phylogenetic analysis and clustering into OTUs

We built an Arcellinida sequence database including sequences derived from identified organisms taken from GenBank as well as from other metabarcoding studies (González-Miguéns et al., 2023; Useros et al., 2024). We aligned the sequences using the auto MAFFT algorithm (Katoh and Standley, 2013), and refined the alignment manually using Geneious Prime (v. 2025.0.3). We constructed a phylogenetic tree in order to cluster all obtained ASVs to operational taxonomic units (OTUs), a proxy for species. We evaluated tree topologies and node supports with Maximum Likelihood (ML), using IQ-TREE2 version 2.0 (Minh et al., 2020), and selected the best substitution models with ModelFinder (Kalyaanamoorthy et al., 2017), implemented in IQ-TREE2 version 2.0. We assessed node supports with 10,000 ultrafast bootstrap replicates approximation (Hoang et al., 2018; Minh et al., 2013), editing the resulting tree in FigTree version 1.4.3. We also generated a patristic distance matrix in Geneious Prime.

We clustered ASV into OTUs, using a distance matrix obtained with Geneious Prime, with the ASV tree as input. We set the cut off (“barcoding gap”) from a molecular pairwise distance of 3% as determined in (Useros et al., 2023) and earlier publications (Kosakyan et al., 2013; Singer et al., 2019). We performed the clustering in R v4.3.2 (R Core Team, 2024) with the DECIPHER (Wright, Erik, 2016) package, using the “complete method”, meaning that the maximum distance between any pair of ASVs from a cluster is 0.03. From the ASV tree, we used the “ape” package (Paradis and Schliep, 2019)to transform the ASV tree into an OTU tree, leaving only one representative per OTU.

### 2.5. Environmental data

To assess the impact of terrestrial land use on freshwater microbial communities, we characterized environmental conditions at each sampling site using four complementary datasets:

#### Land use

We used raster from surrounding land cover from Spanish CORINE Land Cover 2018 maps (Büttner, 2014) and shapefiles of population, population density, agricultural and livestock farms from Ministry for Ecological Transition and Demographic Challenge of Spain and road and rail network from Download Centre of the Spanish National Centre for Geographic Information (https://centrodedescargas.cnig.es/CentroDescargas/home). Subsequently, we converted the Shapefiles into raster images with a resolution of 400 m, applying a 2 km buffer around each point, except for land cover data, which had a 20 m buffer.

#### Soil characteristics

We extracted soil properties e.g., texture, organic matter content and silt from rasters of SoilGrids 2.0 (Poggio et al., 2021) with a resolution of ∼269 m to capture the influence of terrestrial substrate on freshwater microbial habitats.

#### Climate

We obtained site-specific climatic conditions of 1 km resolution raster from CHELSA (Brun et al., 2022; Karger et al., 2017) including mean annual temperature, precipitation, and relevant seasonal variables (i.e. all bioclim variables).

#### Water chemistry

We performed in situ / laboratory measurements on water samples to quantify pH, temperature, conductivity, nitrate, dissolved oxygen, suspended solids, ammonium and the quality state of water bodies following internal protocols by Eurofins Cavendish S.L.U.

We standardized all environmental variables prior to statistical analysis. We examined correlations among variables to reduce redundancy and retained only non-collinear descriptors for downstream analyses linking microbial diversity to land use and habitat conditions (Table S2).

### 2.6. Data analyses

To disentangle how land-atmosphere-water linkages structure Arcellinida communities, we integrated 36 environmental predictors across four categories: regional climate (BIOCLIM+ variables), soil biogeochemistry (ISRIC global soils database), terrestrial land use (agricultural, urban, and land cover), and in-water chemistry (Table S3). We performed a principal component analysis (PCA) to condense these multidimensional gradients into five interpretable axes per category (Fig. S1; Tables S3, S4).

We filtered OTU abundances to retain only those present in more than two samples (presence > 2), and we added total diversity (expressed in Shannon indexes) to the response set. To stabilise variances and reduce the influence of extreme values, we applied a log1p transformation to all response columns (OTUs and diversity; Table S5). Prior to calculation, we excluded samples with fewer than three observed OTUs (i.e., richness ≤ 2) from the diversity analysis to avoid bias due to low community complexity. We calculated Shannon indexes from the OTU abundance table using the “diversity” function from the “vegan” package (Oksanen et al., 2020; v2.6-8) in R. For the excluded samples, the diversity value was set as ‘NA’ to indicate missing data.

We implemented all models in R v4.3.2 using the “caret” (Kuhn, 2008; v7.0-1), “RandomForest” (Breiman, 2001; v4.7-1.2) and “pdp” (Greenwell, Brandon, 2017; v0.8.2) packages. We fitted an independent model for each column of the response matrix (each OTU and Shannon indexes; Table S5). We processed the response variable as numerical (regression) when it had ≥5 distinct values, and as a factor (classification) otherwise. As predictors, we used the five axes of the PCA performed for each of the four categories of environmental variables (land use, climate, soil and water; see above), thus representing a total of 20 variables (Tables S3, S4). We partitioned the data into a training set (75%) and a test set (25%). For each model, we trained 1,000 trees on the training set and optimised the number of variables randomly sampled at each split (mtry) through tuning with repeated 3-fold cross-validation. Before fitting the models, we centred and scaled the predictor variables. We evaluated model performance on the test set using R², root mean squared error (RMSE), and mean absolute error (MAE) for numeric responses, and accuracy and Kappa statistics for categorical responses.

We extracted variable importance for each model using the “varImp” function of “caret” package. To facilitate thematic interpretation, we grouped predictors into categories based on patterns in their variable names: land use, climate, soil, and water. For taxa with model performance R² > 0.1, we compared the distribution of variable importance among categories using ANOVA followed by Tukey post hoc comparisons (TukeyHSD), and we generated boxplots by category.

For each model, we calculated partial dependence plots (PDPs) at the marginal level using the “pdp” package, evaluating the effect of each predictor (first axis of the PCA for each category of environmental variables) on the estimated response (total diversity), while maintaining the other predictors in their observed distribution.

To summarise the direction and intensity of the marginal effects, we calculated correlations between each predictor axis and the response for every OTU × variable combination. We obtained p-values using t-transformation for Pearson correlations (p < 0.05). We visualised these results as a heat map with row clustering using “pheatmap” function of “pheatmap” package (Kolde, 2025).

## 3. Results

### 3.1 Arcellinida diversity across Mediterranean freshwater ecosystems

High-throughput, Arcellinida-specific sequencing of the mitochondrial COI barcode region yielded 8.2 million quality-filtered reads after rigorous curation to remove chimeras, singletons, and tag-jumping artefacts (see *Materials and methods*). These reads resolved into a total of 1,654 amplicon sequence variants (ASVs, corresponding to infraspecific variation (González-Miguéns et al., 2024)) representing 590 operational taxonomic units (OTUs) based on species-level barcoding gaps established for Arcellinida (Fig. S2, Tables S6-S10).

The resulting dataset revealed remarkably high diversity, with 1,171 unique ASVs (71% of all ASVs); community composition varied substantially across waterbodies, with rank abundance curves showing typical ecological patterns of a few dominant taxa and many rare ones (Fig. S2).

### 3.2. Terrestrial landscapes shape aquatic microbial diversity

We integrated five interpretable axes per variable category as environmental predictors (Fig. S1; Tables S2-S4), along with predominant OTU and the total diversity as response to modelled.

Random Forest classification models revealed a hierarchy of cross-ecosystem influence. The models showed predictive ability for 15 OTUs and total diversity (R²: 0.10–0.89, median = 0.45; Table S11), with evaluation metrics ranging from 0.103 to 0.890 R², 0.198–2.462 RMSE and 0.055–0.842 MAE (Table S11). Exploring the variables importance analysis in the model, the importance of each group of variables shows significant differences in land use and water variables (Fig. 2).

**Fig. 2.**
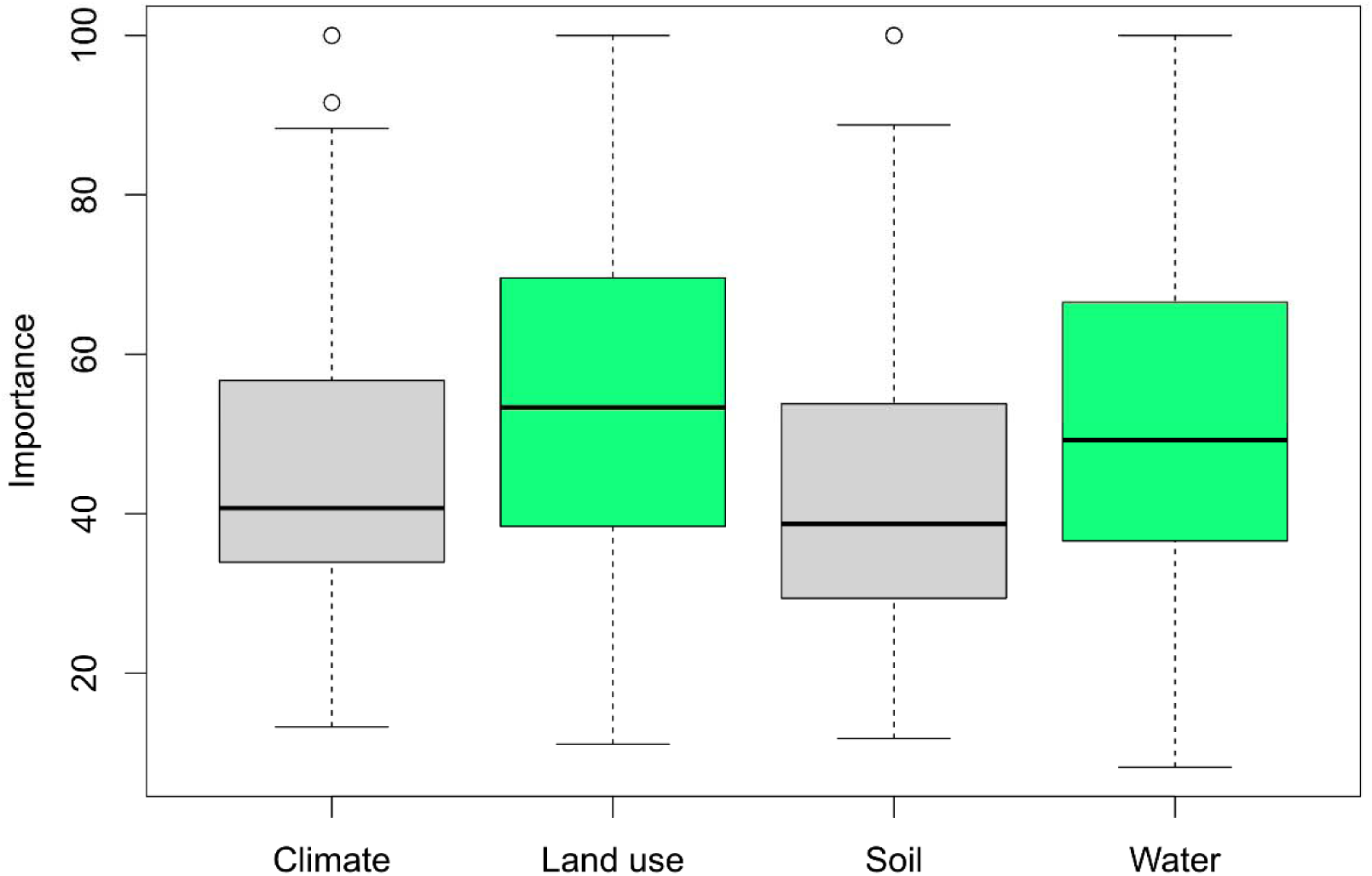
Environmental drivers of Arcellinida diversity. Relative contribution of four environmental categories (land use, climate, soil and water) to observed diversity patterns. Boxplots show statistically significant differences among categories (p < 0.05); boxplots sharing the same colour do not differ significantly.

### 3.3. Response of Arcellinida to land use impacts

Individual taxa exhibited complex and often non-linear responses to terrestrial land use and associated environmental gradients. For example, within genus *Netzelia*, OTU98 declined sharply with land use intensity, whereas OTU102 persisted (Fig. 3; A). In general, such a pattern tends to repeat itself; diversity declined consistently with increasing land use intensity. Total community diversity (Shannon diversity) integrated these idiosyncratic responses into a coherent signal that reflected the cumulative impact of human modification (Fig. 3; A).

**Fig. 3.**
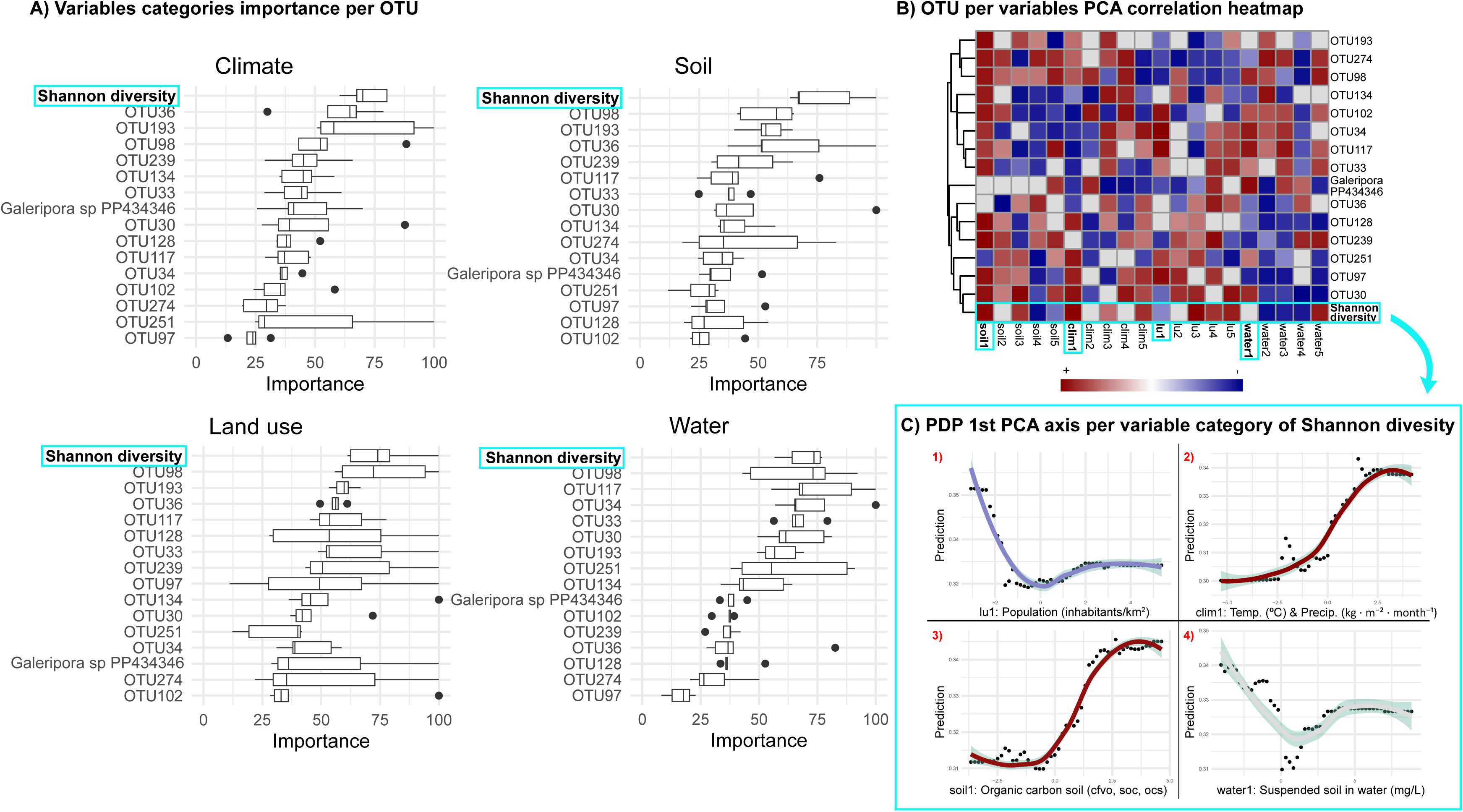
Influence of environmental variables on Arcellinida diversity and distribution. (A) Variable importance analysis showing the relative contribution of four environmental categories (land use, climate, soil, and water) to models explaining total diversity (Shannon index) and individual OTU distributions. (B) Correlation heat map showing relationships between diversity metrics (Shannon index and OTU occurrence) and the first five principal component analysis (PCA) axes derived from all environmental variables. Red indicates positive correlations and blue indicates negative correlations. We used the first PCA axis of each environmental category and total diversity (highlighted in light blue) as predictors for partial dependence plots. Rows (OTUs) are hierarchically clustered using Euclidean distance and complete linkage, grouping taxa with similar responses to environmental gradients. (C) Partial dependence plots showing the relationship between total diversity and the best predictors on the first PCA axis of each environmental category. Curves represent changes in predicted diversity as the focal variable varies, while all other predictors are held constant at their mean values.

Land use correlation heat map showed relationships between Shannon index and the different OTU occurrence and the first five principal component analysis (PCA) axes. Of the 16 OTUs assessed against the five land-use intensity axes (80 OTU-axis pairs in total), 43.75% showed positive correlations, 36.25% showed negative correlations and 20% showed neutral correlations (Fig.3; B).

### 3.4. Terrestrial land use drives aquatic diversity through threshold dynamics

Partial dependence plots revealed that terrestrial human activity exerts non-linear control over aquatic protist communities (Fig. 3; C). Arcellinida diversity peaked in the least populated sites but declined precipitously with increasing human density, stabilizing at a plateau beyond ∼33 inhabitants/km² (Fig. 3; C1). Soil organic carbon, a parameter associated to soil degradation under human influence also increases to a plateau; sites with depleted soil carbon supported less diverse aquatic assemblages (Fig. 3; C3).

Water nutrient states mirrored also this non-linearity: diversity decreased steeply from oligotrophic to eutrophic conditions before plateauing in highly nutrient-enriched systems (Fig. 3; C4).

## 4. Discussion

### 4.1 Arcellinida diversity primarily reflects land use

Freshwater ecosystems are closely linked to their surrounding landscapes, with human activities reshaping both terrestrial and aquatic environments deeply. Our study demonstrates water chemistry and land use act as complementary drivers, with both contributing substantially to observed patterns in Arcellinida diversity, but with land use providing the most consistent broad-scale signal. This hierarchy of impacts reflects a potential cascade of effects: land use alters soil properties, which in turn modify adjacent water conditions, ultimately shaping aquatic biodiversity. Crucially, while physicochemical water parameters fluctuate frequently, land use transforms ecosystems over decadal timescales, leaving lasting imprints on resilient organisms like Arcellinida. Together, these results demonstrate that the reorganization of freshwater microbial communities across the Mediterranean landscape is governed by a coupled climate–land use system. The imprint of human activity—expressed through regional warming, terrestrial carbon loss, and nutrient enrichment—extends from soils to surface waters, emphasizing that conserving aquatic biodiversity in the Anthropocene requires managing both land and climate as a single, integrated continuum.

The significant impact of land use variables in our Random Forest models (Fig. 2) stems from the unique life history traits of Arcellinida. With generation times exceeding 10 days (Lousier, 1974) and their ability to enter dormant states during unfavourable conditions (Ogden and Meisterfeld, 1991), these protists integrate environmental changes over long periods of time, making them exceptionally sensitive to persistent landscape alterations. This temporal decoupling explains why water parameters, while significant, showed weaker correlations with community patterns; their more rapid fluctuations are effectively buffered by Arcellinida biological inertia. Our results align with emerging understanding of how terrestrial processes govern aquatic ecosystems (Dahlin et al., 2021), particularly in Mediterranean regions where land use intensification interacts with increasing aridification (Estrela-Segrelles et al., 2023).

### 4.2 Threshold pattern response to land use and cascading effects

Total diversity responds in a threshold pattern, which illustrates how land-based human activities— agriculture, urbanization, infrastructure—are linked through catchments to restructure aquatic microbiomes. These impacts saturate once landscapes cross critical development thresholds (Fig. 3; C1). In addition, since nutrient loading is the primary aquatic signature of terrestrial land use change, water nutrient pattern confirms that what happens on land—fertilizer application, wastewater discharge, soil erosion—is linked to the structure of freshwater microbial life through biogeochemical cascades. The threshold at intermediate nutrient levels marks a transition beyond which aquatic ecosystems lose their capacity to support diverse protist assemblages (Fig. 3; C4). On the other hand, warmer climates— amplified by ongoing climate change—were associated with higher protist diversity up to intermediate levels, likely by expanding metabolic and habitat niches before thermal stress began to constrain community composition (Fig. 3; C2). Precipitations during the warmest quarter of the year had also a positive effect until an intermediate level, after which summer rainstorms, increasingly destructive under a regional climate change scenario, acted negatively on diversity. A similar pattern was observed for soil composition (SOC), with soils with higher levels of SOC were associated with higher protist diversity (Fig. 3; C3). These relationships reveal how climate and land use operate as intertwined dimensions of human impact: warming is associated with altered hydrological regimes and nutrient fluxes, while land use practices relate to the carbon and energy inputs that sustain aquatic life. Therefore, the influence of human activity in the Anthropocene extends beyond terrestrial boundaries, propagating through interconnected Earth systems to reshape even microscopic aquatic life.

### 4.3 Increasing human land use intensity reduces community complexity by filtering out sensitive taxa

The response of Arcellinida to environmental gradients appears to operate primarily at the community level, where emergent properties integrate signals across multiple taxa. Accordingly, total diversity, as measured by the Shannon index, proved to be more sensitive to environmental gradients than individual taxa alone. Together, these results indicate that Arcellinida-based assessment is most effective when combining community-level diversity metrics with selected taxon-level indicators, rather than relying on a single metric to capture all aspects of environmental change.

This pattern was particularly evident in sites located in agricultural and urban areas, where we observed both a reduction in diversity and early signs of biotic homogenisation (Fig. 3). Nevertheless, individual taxa can still exhibit strong but variable responses to environmental change. In some cases, even closely related species may display contrasting ecological responses. The genus *Netzelia* exemplifies this pattern, with species showing differing sensitivities to environmental gradients (Fig. 3; OTU98 and OTU102). As landscapes shift from natural to managed or urbanized states, local environmental filters tighten: some OTUs (=sensitive specialists) disappear while others (generalists) persist, leading to a measurable erosion of community complexity (Fig. 3; A). This type of species filtering is consistent with broader observations of biodiversity responses to anthropogenic disturbance, including the processes described by McKinney and Lockwood (McKinney and Lockwood, 1999), in which environmental pressures favour tolerant taxa while reducing overall diversity.

The mechanisms underlying these patterns likely involve multiple stressor pathways. Intensive land use reduces soil organic carbon inputs to aquatic systems while increasing pollutant loads (Schürings et al., 2022), creating conditions that favour stress-tolerant generalists over niche specialists. Our partial dependence plots (Fig. 3; C) reinforce this interpretation, showing positive diversity associations with high soil organic carbon (Fig. 3; C3) but negative relationships with eutrophication indicators (Fig. 3; C4) (such as low oxygen and high nitrogen content). These findings extend previous work on protist bioindication (Payne, 2013) by demonstrating how specific taxa reflect particular aspects of catchment health.

From a conservation perspective, our results call for a paradigm shift in freshwater management. Traditional water quality control is based on physicochemical parameters and aquatic taxa monitoring (Horton R. K., 1965). While these studies remain important, they fail to capture the processes at the catchment scale that ultimately determine ecosystem health. Instead, we advocate for integrated water and land management that recognises: (1) the primacy of terrestrial processes in shaping aquatic biodiversity, (2) the value of microbial indicators such as Arcellinida in the early detection of ecosystem degradation, and (3) the need to monitor specialised taxa as sentinels of ecological integrity.

Our study establishes the importance of land use variables in shaping freshwater protist diversity, yet captures a single temporal snapshot during one seasonal period; multi-seasonal and inter-annual sampling would clarify how these patterns vary across Mediterranean hydrological cycles. Future work should combine experimental manipulations, broader taxonomic coverage, global mapping of Arcellinida distribution, and multi-regional comparisons to validate causal mechanisms and develop standardized Arcellinida-based bioassessment protocols applicable across diverse Mediterranean freshwater systems.

## 5. Conclusions

By bridging landscape ecology and protist biodiversity, we demonstrate how human activities reshape aquatic communities through terrestrial pathways, a perspective that is fundamental to ecosystem conservation. Given that Mediterranean freshwater systems face increasing threats from climate change and intensified land use, incorporating protist indicators into monitoring programmes could provide the early warning system needed to prevent irreversible biodiversity loss as well as human life quality. Ultimately, recognising the terrestrial imprint on aquatic microbial biodiversity will be key to conserving freshwater resilience under accelerating global change.

## CRediT authorship contribution statement

**Carmen Soler-Zamora**: Conceptualization, Data curation, Formal analysis, Funding acquisition, Investigation, Methodology, Project administration, Resources, Software, Visualization, Writing – original draft.

**Emilio Cano**: Data curation, Methodology.

**Patrizia Elena Vannucchi**: Funding acquisition, Methodology, Resources, Visualization.

**Enrique Lara**: Conceptualization, Funding acquisition, Methodology, Supervision, Validation, Writing – original draft.

**Bertrand Fournier**: Conceptualization, Formal analysis, Methodology, Software, Supervision, Validation, Writing – original draft.

## Acknowledgements

This publication is part of the project “Research and development of a novel biological water quality index based on Arcellinida using metabarcoding techniques” from Eurofins Cavendish company, funded by the CDTI (Centre for Industrial Technological Development), supported by the Spanish Ministry of Science and Innovation, under the R&D projects (Grant number IDI-20220885). The collaboration necessary for the completion of this publication was made possible thanks to CSIC iMOVE’s support for short stays at R&D&I centres awarded to C.S.Z. (Grant number IMOVE24036). E. L. was supported by a grant from the Spanish Government (Programa Generación de Conocimiento), grant number PID2021-128499NB-I00, 10.13039/501100011033, (MCIU/AEI/ FEDER, UE). The authors would like to thank the Aquatic Ecology Department at Eurofins Cavendish for their valuable support and collaboration throughout the fieldwork.

## Data availability

We deposited the Illumina raw DNA sequences in GenBank BioProject PRJNA1454412. All raw data, models, code, statistical analyses and supplementary tables and figures used in this study can be found on Zenodo (https://doi.org/10.5281/zenodo.19885362).

## References

Albert, J.S., Destouni, G., Duke-Sylvester, S.M., Magurran, A.E., Oberdorff, T., Reis, R.E., Winemiller, K.O., Ripple, W.J., 2021. Scientists’ warning to humanity on the freshwater biodiversity crisis. Ambio 50, 85–94. 10.1007/s13280-020-01318-8

Allan, J.D., Flecker, A.S., 1993. Biodiversity Conservation in Running Waters. Bioscience 43, 32–43. 10.2307/1312104

Bjorbækmo, M.F.M., Evenstad, A., Røsæg, L.L., Krabberød, A.K., Logares, R., 2020. The planktonic protist interactome: where do we stand after a century of research? ISME J. 14, 544–559. 10.1038/s41396-019-0542-5

Breiman, L., 2001. Random Forests. Mach. Learn. 45, 5–32. 10.1023/A:1010933404324

Brun, P., Zimmermann, N.E., Hari, C., Pellissier, L., Karger, D., 2022. CHELSA-BIOCLIM+ A novel set of global climate-related predictors at kilometre-resolution. EnviDat. https://www.doi.org/10.16904/envidat.332.

Büttner, G., 2014. CORINE Land Cover and Land Cover Change Products, in: Remote Sensing and Digital Image Processing. pp. 55–74. 10.1007/978-94-007-7969-3_5

Callahan, B.J., McMurdie, P.J., Rosen, M.J., Han, A.W., Johnson, A.J.A., Holmes, S.P., 2016. DADA2: High-resolution sample inference from Illumina amplicon data. Nat. Methods 13, 581–583. 10.1038/nmeth.3869

Carpenter, S.R., Caraco, N.F., Correll, D.L., Howarth, R.W., Sharpley, A.N., Smith, V.H., 1998. Nonpoint pollution of surface waters with phosphorus and nitrogen. Ecological Applications 8, 559–568. 10.1890/1051-0761(1998)008[0559:NPOSWW]2.0.CO;2

Charqueño Celis, N.F., Garibay, M., Sigala, I., Brenner, M., Echeverria-Galindo, P., Lozano, S., Massaferro, J., Pérez, L., 2019. Testate amoebae (Amoebozoa: Arcellinidae) as indicators of dissolved oxygen concentration and water depth in lakes of the Lacandón Forest, southern Mexico. J. Limnol. 79, 82–91. 10.4081/jlimnol.2019.1936

Dahlin, K.M., Zarnetske, P.L., Read, Q.D., Twardochleb, L.A., Kamoske, A.G., Cheruvelil, K.S., Soranno, P.A., 2021. Linking Terrestrial and Aquatic Biodiversity to Ecosystem Function Across Scales, Trophic Levels, and Realms. Front. Environ. Sci. 9. 10.3389/fenvs.2021.692401

ElKhouri-Vidarte, N., Useros, F., Lara, E., 2025. Beneath the Cedars: Exploring the Water-Energy Balance on Arcellinida Biodiversity in Lebanon’s Cedar Forests. Microb. Ecol. 89, 33. 10.1007/s00248-025-02666-2

Escobar, J., Brenner, M., Whitmore, T.J., Kenney, W.F., Curtis, J.H., 2008. Ecology of testate amoebae (thecamoebians) in subtropical Florida lakes. J. Paleolimnol. 40, 715–731. 10.1007/s10933-008-9195-5

Estrela-Segrelles, C., Gómez-Martínez, G., Pérez-Martín, M.Á., 2023. Climate Change Risks on Mediterranean River Ecosystems and Adaptation Measures (Spain). Water Resources Management 37, 2757–2770. 10.1007/s11269-023-03469-1

Flinn, K.M., Vellend, M., 2005. Recovery of forest plant communities in post-agricultural landscapes. Front. Ecol. Environ. 10.1890/1540-9295(2005)003[0243:ROFPCI]2.0.CO;2

Folmer, O., Black, M., Hoeh, W., Lutz, R., Vrijenhoek, R., 1994. DNA primers for amplification of mitochondrial cytochrome c oxidase subunit I from diverse metazoan invertebrates. Mol. Mar. Biol. Biotechnol. 3, 294–299.

Fournier, B., Samaritani, E., Frey, B., Seppey, C.V.W., Lara, E., Heger, T.J., Mitchell, E.A.D., 2020. Higher spatial than seasonal variation in floodplain soil eukaryotic microbial communities. Soil Biol. Biochem. 147, 107842. 10.1016/j.soilbio.2020.107842

González-Miguéns, R., Cano, E., García-Gallo Pinto, M., Peña, P.G., Rincón-Barrado, M., Iglesias, G., Blanco-Rotea, A., Carrasco-Braganza, M.I., de Salvador-Velasco, D., Guillén-Oterino, A., Tenorio-Rodríguez, D., Siemensma, F., Velázquez, D., Lara, E., 2024. The voice of the little giants: Arcellinida testate amoebae in environmental DNA-based bioindication, from taxonomy free to haplotypic level. Mol. Ecol. Resour. 24, e13999. 10.1111/1755-0998.13999

González-Miguéns, R., Cano, E., Guillén-Oterino, A., Quesada, A., Lahr, D.J.G., Tenorio-Rodríguez, D., de Salvador-Velasco, D., Velázquez, D., Carrasco-Braganza, M.I., Patterson, R.T., Lara, E., Singer, D., 2023. A needle in a haystack: A new metabarcoding approach to survey diversity at the species level of Arcellinida (Amoebozoa: Tubulinea). Mol. Ecol. Resour. 23, 1034–1049. 10.1111/1755-0998.13771

González-Miguéns, R., Gàlvez-Morante, À., Skamnelou, M., Antó, M., Casacuberta, E., Richter, D.J., Lara, E., Vaulot, D., Del Campo, J., Ruiz-Trillo, I., 2025. A novel taxonomic database for eukaryotic mitochondrial cytochrome oxidase subunit I gene (eKOI), with a focus on protists diversity. Database 2025. 10.1093/DATABASE/BAAF057

González-Miguéns, R., Soler-Zamora, C., Villar-Depablo, M., Todorov, M., Lara, E., 2022. Multiple convergences in the evolutionary history of the testate amoeba family Arcellidae (Amoebozoa: Arcellinida: Sphaerothecina): when the ecology rules the morphology. Zool. J. Linn. Soc. 194, 1044– 1071. 10.1093/zoolinnean/zlab074

Greenwell, Brandon, M., 2017. pdp: An R Package for Constructing Partial Dependence Plots. R J. 9, 421. 10.32614/RJ-2017-016

Guo, P., Li, C., Liu, J., Chai, B., 2023. Predation has a significant impact on the complexity and stability of microbial food webs in subalpine lakes. Microbiol. Spectr. 11. 10.1128/spectrum.02411-23

Haddad, N.M., Brudvig, L.A., Clobert, J., Davies, K.F., Gonzalez, A., Holt, R.D., Lovejoy, T.E., Sexton, J.O., Austin, M.P., Collins, C.D., Cook, W.M., Damschen, E.I., Ewers, R.M., Foster, B.L., Jenkins, C.N., King, A.J., Laurance, W.F., Levey, D.J., Margules, C.R., Melbourne, B.A., Nicholls, A.O., Orrock, J.L., Song, D.-X., Townshend, J.R., 2015. Habitat fragmentation and its lasting impact on Earth’s ecosystems. Sci. Adv. 1. 10.1126/sciadv.1500052

Hoang, D.T., Chernomor, O., von Haeseler, A., Minh, B.Q., Vinh, L.S., 2018. UFBoot2: Improving the Ultrafast Bootstrap Approximation. Mol. Biol. Evol. 35, 518–522. 10.1093/molbev/msx281

Horton R. K., 1965. An index number system for rating water quality. Journal of the Water Pollution Control Federation 37, 300–306.

Kalyaanamoorthy, S., Minh, B.Q., Wong, T.K.F., von Haeseler, A., Jermiin, L.S., 2017. ModelFinder: fast model selection for accurate phylogenetic estimates. Nat. Methods 14, 587–589. 10.1038/nmeth.4285

Karger, D.N., Conrad, O., Böhner, J., Kawohl, T., Kreft, H., Soria-Auza, R.W., Zimmermann, N.E., Linder, H.P., Kessler, M., 2017. Climatologies at high resolution for the earth’s land surface areas. Sci. Data 4, 170122. 10.1038/sdata.2017.122

Katoh, K., Standley, D.M., 2013. MAFFT Multiple Sequence Alignment Software Version 7: Improvements in Performance and Usability. Mol. Biol. Evol. 30, 772–780. 10.1093/molbev/mst010

Kolde, R., 2025. pheatmap: Pretty Heatmaps. CRAN: Contributed Packages. 10.32614/CRAN.package.pheatmap

Kosakyan, A., Gomaa, F., Mitchell, E.A.D., Heger, T.J., Lara, E., 2013. Using DNA-barcoding for sorting out protist species complexes: A case study of the Nebela tincta–collaris–bohemica group (Amoebozoa; Arcellinida, Hyalospheniidae). Eur. J. Protistol. 49, 222–237. 10.1016/j.ejop.2012.08.006

Kosakyan, A., Lara, E., 2019. Using Testate Amoebae Communities to Evaluate Environmental Stress: A Molecular Biology Perspective, in: Encyclopedia of Environmental Health. Elsevier, pp. 308–313. 10.1016/B978-0-12-409548-9.11589-1

Kuhn, M., 2008. Building Predictive Models in R Using the caret Package. J. Stat. Softw. 28, 1–26. 10.18637/jss.v028.i05

Lousier, J.D., 1974. Effects of experimental soil moisture fluctuations on turnover rates of testacea. Soil Biol. Biochem. 6, 19–26. 10.1016/0038-0717(74)90006-6

Lynch, A.J., Cooke, S.J., Arthington, A.H., Baigun, C., Bossenbroek, L., Dickens, C., Harrison, I., Kimirei, I., Langhans, S.D., Murchie, K.J., Olden, J.D., Ormerod, S.J., Owuor, M., Raghavan, R., Samways, M.J., Schinegger, R., Sharma, S., Tachamo-Shah, R., Tickner, D., Tweddle, D., Young, N., Jähnig, S.C., 2023. People need freshwater biodiversity. WIREs Water 10. 10.1002/wat2.1633

McFadden, I.R., Sendek, A., Brosse, M., Bach, P.M., Baity-Jesi, M., Bolliger, J., Bollmann, K., Brockerhoff, E.G., Donati, G., Gebert, F., Ghosh, S., Ho, H., Khaliq, I., Lever, J.J., Logar, I., Moor, H., Odermatt, D., Pellissier, L., de Queiroz, L.J., Rixen, C., Schuwirth, N., Shipley, J.R., Twining, C.W., Vitasse, Y., Vorburger, C., Wong, M.K.L., Zimmermann, N.E., Seehausen, O., Gossner, M.M., Matthews, B., Graham, C.H., Altermatt, F., Narwani, A., 2023. Linking human impacts to community processes in terrestrial and freshwater ecosystems. Ecol. Lett. 26, 203–218. 10.1111/ele.14153

McKinney, M.L., Lockwood, J.L., 1999. Biotic homogenization: a few winners replacing many losers in the next mass extinction. Trends Ecol. Evol. 14, 450–453. 10.1016/S0169-5347(99)01679-1

Minh, B.Q., Nguyen, M.A.T., von Haeseler, A., 2013. Ultrafast Approximation for Phylogenetic Bootstrap. Mol. Biol. Evol. 30, 1188–1195. 10.1093/molbev/mst024

Minh, B.Q., Schmidt, H.A., Chernomor, O., Schrempf, D., Woodhams, M.D., von Haeseler, A., Lanfear, R., 2020. IQ-TREE 2: New Models and Efficient Methods for Phylogenetic Inference in the Genomic Era. Mol. Biol. Evol. 37, 1530–1534. 10.1093/molbev/msaa015

Nasser, N.A., Patterson, R.T., Roe, H.M., Galloway, J.M., Falck, H., Sanei, H., 2020. Use of Arcellinida (testate lobose amoebae) arsenic tolerance limits as a novel tool for biomonitoring arsenic contamination in lakes. Ecol. Indic. 113, 106177. 10.1016/j.ecolind.2020.106177

Ogden, C.G., Meisterfeld, R., 1991. The biology and ultrastructure of the testate amoeba, Difflugia lucida penard (Protozoa, Rhizopoda). Eur. J. Protistol. 26, 256–269. 10.1016/S0932-4739(11)80147-1

Oksanen, J., Simpson, G.L., Blanchet, F.G., Kindt, R., Legendre, P., Minchin, P.R., O’Hara, R.B., Solymos, P., Stevens, M.H.H., Szoecs, E., Wagner, H., Bedward, M., Bolker, B., Borcard, D., Carvalho, G., De Caceres, M., Durand, S., Evangelista, H.B.A., Hannigan, G., Hill, M.O., Lahti, L., Martino, C., Ouellette, M.-H., Ribeiro Cunha, E., Smith, T., Stier, A., Ter Braak, C.J.F., Weedon, J., 2026. vegan: Community Ecology Package. R package version 2.6–8. 10.32614/CRAN.package.vegan

Paradis, E., Schliep, K., 2019. ape 5.0: an environment for modern phylogenetics and evolutionary analyses in R. Bioinformatics 35, 526–528. 10.1093/bioinformatics/bty633

Patterson, R.T., Lamoureux, E.D.R., Neville, L.A., Macumber, A.L., 2013. Arcellacea (Testate Lobose Amoebae) as pH Indicators in a Pyrite Mine-Acidified Lake, Northeastern Ontario, Canada. Microb. Ecol. 65, 541–554. 10.1007/s00248-012-0108-9

Payne, R.J., 2013. Seven reasons why protists make useful bioindicators. Acta Protozool. 52, 105–113. 10.4467/16890027AP.13.0011.1108

Peay, K.G., Garbelotto, M., Bruns, T.D., 2010. Evidence of dispersal limitation in soil microorganisms: Isolation reduces species richness on mycorrhizal tree islands. Ecology 91, 3631–3640. 10.1890/09-2237.1

Penard, E., 1902. Faune rhizopodique du bassin du Léman. H. Kündig, Genève. 10.5962/bhl.title.1711

Poggio, L., De Sousa, L.M., Batjes, N.H., Heuvelink, G.B.M., Kempen, B., Ribeiro, E., Rossiter, D., 2021. SoilGrids 2.0: Producing soil information for the globe with quantified spatial uncertainty. SOIL 7, 217–240. 10.5194/soil-7-217-2021

R Core Team, 2024. R: A language and environment for statistical computing. R Foundation for Statistical Computing, Vienna, Austria.

Reinhardt, E.G., Dalby, A.P., Kumar, A., Patterson, R.T., 1998. Arcellaceans as Pollution Indicators in Mine Tailing Contaminated Lakes near Cobalt, Ontario, Canada. Micropaleontology 44, 131. 10.2307/1486066

Rodriguez-Martinez, S., Klaminder, J., Morlock, M.A., Dalén, L., Huang, D.Y., 2023. The topological nature of tag jumping in environmental DNA metabarcoding studies. Mol. Ecol. Resour. 23, 621–631. 10.1111/1755-0998.13745

Roe, H.M., Patterson, R.T., Swindles, G.T., 2010. Controls on the contemporary distribution of lake thecamoebians (testate amoebae) within the Greater Toronto Area and their potential as water quality indicators. J. Paleolimnol. 43, 955–975. 10.1007/s10933-009-9380-1

Rognes, T., Flouri, T., Nichols, B., Quince, C., Mahé, F., 2016. VSEARCH: a versatile open source tool for metagenomics. PeerJ 4, e2584. 10.7717/peerj.2584

Sayer, C.A., Fernando, E., Jimenez, R.R., Macfarlane, N.B.W., Rapacciuolo, G., Böhm, M., Brooks, T.M., Contreras-MacBeath, T., Cox, N.A., Harrison, I., Hoffmann, M., Jenkins, R., Smith, K.G., Vié, J.-C., Abbott, J.C., Allen, D.J., Allen, G.R., Barrios, V., Boudot, J.-P., Carrizo, S.F., Charvet, P., Clausnitzer, V., Congiu, L., Crandall, K.A., Cumberlidge, N., Cuttelod, A., Dalton, J., Daniels, A.G., De Grave, S., De Knijf, G., Dijkstra, K.-D.B., Dow, R.A., Freyhof, J., García, N., Gessner, J., Getahun, A., Gibson, C., Gollock, M.J., Grant, M.I., Groom, A.E.R., Hammer, M.P., Hammerson, G.A., Hilton-Taylor, C., Hodgkinson, L., Holland, R.A., Jabado, R.W., Juffe Bignoli, D., Kalkman, V.J., Karimov, B.K., Kipping, J., Kottelat, M., Lalèyè, P.A., Larson, H.K., Lintermans, M., Lozano, F., Ludwig, A., Lyons, T.J., Máiz-Tomé, L., Molur, S., Ng, H.H., Numa, C., Palmer-Newton, A.F., Pike, C., Pippard, H.E., Polaz, C.N.M., Pollock, C.M., Raghavan, R., Rand, P.S., Ravelomanana, T., Reis, R.E., Rigby, C.L., Scott, J.A., Skelton, P.H., Sloat, M.R., Snoeks, J., Stiassny, M.L.J., Tan, H.H., Taniguchi, Y., Thorstad, E.B., Tognelli, M.F., Torres, A.G., Torres, Y., Tweddle, D., Watanabe, K., Westrip, J.R.S., Wright, E.G.E., Zhang, E., Darwall, W.R.T., 2025. One-quarter of freshwater fauna threatened with extinction. Nature 638, 138–145. 10.1038/s41586-024-08375-z

Schnell, I.B., Bohmann, K., Gilbert, M.T.P., 2015. Tag jumps illuminated – reducing sequence-to-sample misidentifications in metabarcoding studies. Mol. Ecol. Resour. 15, 1289–1303. 10.1111/1755-0998.12402

Schürings, C., Feld, C.K., Kail, J., Hering, D., 2022. Effects of agricultural land use on river biota: a meta-analysis. Environ. Sci. Eur. 34, 124. 10.1186/s12302-022-00706-z

Singer, D., Mitchell, E.A.D., Payne, R.J., Blandenier, Q., Duckert, C., Fernández, L.D., Fournier, B., Hernández, C.E., Granath, G., Rydin, H., Bragazza, L., Koronatova, N.G., Goia, I., Harris, L.I., Kajukało, K., Kosakyan, A., Lamentowicz, M., Kosykh, N.P., Vellak, K., Lara, E., 2019. Dispersal limitations and historical factors determine the biogeography of specialized terrestrial protists. Mol. Ecol. 28, 3089–3100. 10.1111/mec.15117

Strayer, D.L., Dudgeon, D., 2010. Freshwater biodiversity conservation: recent progress and future challenges. J. North Am. Benthol. Soc. 29, 344–358. 10.1899/08-171.1

Sun, W., Xia, C., Xu, M., Guo, J., Sun, G., 2017. Seasonality Affects the Diversity and Composition of Bacterioplankton Communities in Dongjiang River, a Drinking Water Source of Hong Kong. Front. Microbiol. 8. 10.3389/fmicb.2017.01644

Townsend, C.R., Uhlmann, S.S., Matthaei, C.D., 2008. Individual and combined responses of stream ecosystems to multiple stressors. Journal of Applied Ecology 45, 1810–1819. 10.1111/j.1365-2664.2008.01548.x

Useros, F., García-Cunchillos, I., Henry, N., Berney, C., Lara, E., 2024. How good are global DNA-based environmental surveys for detecting all protist diversity? Arcellinida as an example of biased representation. Environ. Microbiol. 26. 10.1111/1462-2920.16606

Useros, F., González-Miguéns, R., Soler-Zamora, C., Lara, E., 2023. When ecological transitions are not so infrequent: independent colonizations of athalassohaline water bodies by Arcellidae (Arcellinida; Amoebozoa), with descriptions of four new species. FEMS Microbiol. Ecol. 99, 1–15. 10.1093/femsec/fiad076

Wright, Erik, S., 2016. Using DECIPHER v2.0 to Analyze Big Biological Sequence Data in R. R J. 8, 352. 10.32614/RJ-2016-025

